# Directed Shortest Walk on Temporal Graphs

**DOI:** 10.1101/2022.07.08.499368

**Authors:** Alex Khodaverdian, Nir Yosef

## Abstract

**Background:** The use of graphs as a way of abstracting and representing biological systems has provided a powerful analysis paradigm. Specifically, graph optimization algorithms are routinely used to address various connectivity queries, such as finding paths between proteins in a protein-protein interaction network, while maximizing objectives such as parsimony. While present studies in this field mostly concern static graphs, new types of data now motivate the need to account for changes that might occur to the elements (nodes) that are represented by the graph on the relationships (edges) between them.

**Results and Discussion:** We define the notion of Directed Temporal Graphs as a series of directed subgraphs of an underlying graph, ordered by time, where only a subset of vertices and edges are present. We then build up towards the Time Conditioned Shortest Walk problem on Directed Temporal Graphs: given a series of time ordered directed graphs, find the shortest walk from any given source node at time point 1 to a target node at time *T* ≥ 1, such that the walk is consistent (monotonically increasing) with the timing of nodes and edges. We show, contrary to the Directed Shortest Walk problem which can be solved in polynomial time, that the Time Conditioned Shortest Walk (TCSW) problem is NP-Hard, and is hard to approximate to factor 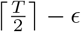 for *T* ≥ 3 and *ε* > 0. Lastly, we develop an integer linear program to solve a generalized version of TCSW, and demonstrate its ability to reach optimality with instances of the human protein interaction network.

**Conclusion:** We demonstrate that when extending the shortest walk problem in computational biology to account for multiple ordered conditions, the problem not only becomes hard to solve, but hard to approximate, a limitation which we address via a new solver. From this narrow definition of TCSW, we relax the constraint of time consistency within the shortest walk, deriving a direct relationship between hardness of approximation and the allowable step size in our walk between time conditioned networks. Lastly we briefly explore a variety of alternative formulations for this problem, providing insight into both tractable and intractable variants.

**Availability:** Our solver for the general k-Time Condition Shortest Walk problem is available at https://github.com/YosefLab/temporal_condition_shortest_walk

## Background

A common approach for investigating biological systems is to first abstract the system as a network. These networks may be defined over broad and varying categories such as species or at a higher resolution over proteins or metabolites within cells. For example, proteins and their pairwise interactions can be modeled as protein protein interaction networks, where nodes represent proteins and edges represent phsyical binding or joint chemical reactions. Once such an abstraction has been made, an investigator can dig into more specific questions. For example, cells express a subset of protein receptors depending on the cell type and cell state. Once a receptor is stimulated via extracellular ligands, a signalling cascade begins at receptor on the cell surface and often ends by modifying several transcription factors in the nucleus. Although a broad range of known pairwise protein interactions are well studied in the protein-protein interaction network, the path in which the signal propagates through this network from a receptor or protein to a terminal protein or transcription factor is less well studied and is a newer question of interest [1, 2, 3, 4, 5]. Under the assumptions of a static protein-protein interaction network, one could simply apply out of the box algorithms such as Djikstra’s algorithm or Steiner Tree to find the most likely path taken [6, 7]. In practice however it turns out that the network is dynamic with time, as only a subset of proteins are active signalling candidates at any given time, be it that the protein may not be present in the cell, or may be deactivated via post-translational modifications [8, 9, 10]. Therefore, the question of interest becomes to not only infer the sequence of protein protein interactions in which receptor signaling leads to changes in transcription factors, but how this signal propagates through a dynamic temporal network.

Temporal networks have been well studied, with applications to broad fields such as communication propagation [11, 12, 13]. The simplest of such temporal problems is the shortest path problem, with the following setup - a network is provided with weighted edges, which travel from timepoint *t*_1_ to *t*_2_, where *t*_2_ > *t*_1_. The goal becomes to find a path which optimizes over a constraint, and is time-monotonic (or increasing in time). Many such possible variants of time-monotonic shortest path exist over this network, for example the earliest arrival time from *a* to *b*, the latest departure time from *a* to *b*, or just simply the lowest cost path (by edge weight), all of which can be solved in polynomial time [14, 15, 16]. There also exist more difficult problems, such as finding strongly connected components [17], or finding time-monotonic spanning trees [18, 19, 20], both of which are intractable, although for the case of minimum spanning trees, there exists an approximation preserving reduction to Directed Steiner Tree, which can be non-trivially approximated [21, 22]. While these works underscore the importance of a singular optimization problem spanning multiple timepoints, they fail to consider several key points present in real biological applications. Firstly, in the aforementioned literature, nodes are always present and traversable. However, in practice, nodes, or proteins, can be in active forms at certain times, and inactive forms at other times. Secondly, a key notion in biology comes from the idea of Occam’s Razor. That is we wish to explain a biological process with the least number edges or nodes traversed, accounting for the fact that the same edge or node can be used multiple times under different contexts (ex. time). This model fails to allow for that possibility due to only allowing for a path like singular edge traversal, rather than a walk.

Another work closely related to this problem comes from Wu et al, who explored the notion of Condition networks [23]. In this setting, a series of directed networks *G_c_* are provided, with the same set of nodes *V*, but a different set of edges *E_c_*. In addition, each network comes with its own set of pairwise connectivity demands *X_c_* = {(*a*_1_, *b*_1_),…, (*a_n_*, *b_n_*)}. The goal becomes to identify a subgraph *H* s.t. all connectivity demands in *X_c_* are satisfied in *H* ⋂ *G_c_*, and *H* is of minimum cost. A special version of this problem is Condition Shortest Path (CSP), in which a singular connectivity demand *X* = (*a, b*) is given for each condition. Both of these problems turn out to be NP-hard to approximate beyond a trivial factor |*C*|. Although this problem attempts to optimize for the shortest path between a given (*a, b*) pair per condition, with information reuse, we are instead interested in a very different scenario. In particular, the scenario we are interested in considers one singular network demand spanning over many graphs, motivated by dynamic biological networks in practice.

## Summary of main contributions

To this end, we introduce the Time Conditioned Shortest Walk (TCSW) problem, which takes on a similar flavor as Condition Shortest Path (CSP) introduced by Wu et al. In this setting we are given a series of ordered networks *G_t_* and ordered conditions {1,…,*T*} representing a discrete measurement of time, and as well as a pair of nodes (*a* ∈ *G*_1_, *b* ∈ *G_T_*). For each time condition, our defined networks *G_t_* have vertices *V_t_* which are allowed to change across networks, but a set of global edges *E* which remain constant. The goal is to find a walk from a source node *a* to target node *b* which begins in *G*_1_, ends in *G_T_*, and satisfies the following constraints: transitions are allowed from *v* ∈ *G_i_* to *w* ∈ *G_i_* or *G*_*i*+1_ if (*v,w*) ∈ *E*. In addition, transitions are allowed from *v* ∈ *G_i_* to *v* ∈ *G*_*i*+1_. Edge costs are only paid once globally; that is, if you reuse an edge, you do not pay a cost.

We first prove a simple algorithm for solving the case for *T* = 2 time conditions. We then move to the more difficult case where vertices are shifting, and show that it is NP-hard to find a solution that achieves an approximation factor better than 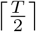 for *T* ≥ 3 via approximation preserving reduction to CSP. We then extend our results to the general setting, where the time differential between two nodes in a path can be an integer *k* greater than 1. We prove that this problem is similarly hard to approximate to a factor better than 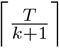, for an arbitrary step size *k*. Lastly, we provide a integer linear program for the general TCSW problem, and show that when provided with real-world input it is capable of finding optimal solutions in reasonable time.

## Definitions and Preliminaries

In graph theory, the shortest path problem is well studied in its many variants: from undirected networks, to weighted directed networks with nonnegative edge weights, to weighted directed networks without negative cycles. Each of these variants can be solved efficiently. For example the most common variant of the shortest path problem over weighted directed networks is solvable via Djikstra’s algorithm in *O*(*E* + *V* log(*V*)) complexity, or with more advanced techniques such as Thorup’s algorithm in time *O*(*E* + *V* log(log(*V*))).

In this paper we extend the generalization of network problems begun under Wu et al to the condition setting. Specifically, we generalize the shortest path problem to the time conditioned setting. Recall that in this setting we have a series of ordered time conditions [*T*] = {1,…,*T*}, each with a corresponding network *G_t_*.

### Definition 1

(Time Condition Shortest Walk (TCSW))
*Given the following inputs:*

1. *A series of directed networks* {*G_t_* = (*V_t_*, *E*)}_*t*∈[*T*]_ *each corresponding to a time condition t. Edges are positively weighted. Note that we denote V* = ∪_*t*_ *V_t_*. *We denote a node v* ∈ *V_T_ as v_t_*
2. *A pair of nodes* (*a*_1_, *b_T_*) *s.t. a*_1_ ∈ *V*_1_ *and b_T_* ∈ *V_T_ which we wish to connect via a walk through G*_1_,…,*G_T_*

Our goal is to find an *a* – *b* walk from *G*_1_ to *G_T_*. We denote a walk *W* in the form *W* = {…, (*v_i_*, *W_j_*),…}, where (*v_i_*, *W_j_*) is a valid edge if *v* ∈ *V_i_*, *w* ∈ *V_j_* and *i* = *j* or *i* = *j* + 1, and (*v,w*) ∈ *E* or *v* = *w*. We denote this “jump” constraint from *G_i_* to *G*_*i*+1_ as the *temporal walk constraint.* The walk begins at *a*_1_ and ends at *b_T_*.

Equivalently, one can consider TCSW over a singular global network defined over all *G_i_*. We define such network 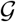 with 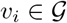 if *v* ∈ *V_i_*, where *v* may be present in multiple *V*\*. An edge *e* = (*v_i_*, *W_j_*) exists in 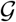 if *v_i_* ∈ *V_i_*, *w_j_* ∈ *V_j_*, *i* = *j* or *i* = *j* + 1, and (*v_i_*, *W_j_*) ∈ *E*. In addition, we add zero weight edges for all nodes *v_i_*, *v*_*i*+1_ if *v_i_* ∈ *V_i_* and *v*_*i*+1_ ∈ *V*_*i*+1_. The goal is to find the minimum cost path from *a*_1_ to *b_T_* where an edge (*v, w*) is only paid for once regardless of however many (*v_i_*, *w_j_*) pairs are traversed.

We offer the most formal presentation of TCSW, its variants, and 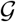 in the Appendix.

#### Definition 2

(*k*–Time Condition Shortest Walk (*k*–TCSW)) *In this variant, we relax the temporal walk constraint to allow for jumps between net-works that aren’t immediately subsequent in time. More specifically: For a given walk W in the form W* = {…,(*v_i_*, *w_j_*),…}, (*v_i_*, *w_j_*) *is a valid edge if* v ∈ *V_i_*, *w* ∈ *V_j_ and i* ≤ *j* ≤ *j* + *k. We denote this relaxed constraint as the k-temporal walk constraint. We note that in the definition for vanilla TCSW we operate under the 1-temporal walk constraint.*

#### Definition 3

(Directed Condition Shortest Path (CSP)) *We draw this definition from Wu et al.*

1. *A sequence of directed graphs G*_1_ = (*V*_1_, *E*), *G*_2_ = (*V*_2_, *E*),…, *G_T_* = (*V_T_*, *E*) *with positively weighted edges and V* = ∪_*t*_ *V_t_*.
2. *A set of C connectivity demands* 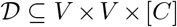 *in the form* (*a*, *b*, 1),…, (*a*, *b*, *C*).

Let *G* = (*V*, *E*) be the underlying network. The goal of this problem is to find a subgraph *H* ⊆ *G* of minimal cost s.t. there exists a path from (*a, b*) in *H* amongst vertices that are active in *G_c_* for all *c*. We note that CSP is hard to approximate to a factor of *C* – *ε* for every *C* ≥ 2 and *ε* > 0, a fact that we will exploit to prove *TCSW* is hard to approximate.

### Problem Variants

The description given above for *TCSW* is just one possible way to describe the Temporal Walk problem. Here we describe several other variants, leaving their analysis for later in the manuscript.

#### Definition 4

(Strict Step Repay – Time Condition Shortest Walk (SSR–TCSW)) *Given the same inputs as TCSW, a demand pair and a series of networks G*_1_…*G_T_*, *the goal is to find an a* – *b walk W from G*_1_ *to G_T_, where W in the form W* ={…, (*v_i_*, *w_j_*),…}, *where* (*v_i_, w_j_*) *is a valid edge if v* ∈ *V_i_, w* ∈ *V_j_* and *i* = *j* + 1, *and* (*v, w*) ∈ *E or v* = *w. That is, every step must move up in time. In addition, edges are paid for per use, i.e. edges can be reused but must be paid for each time*.

#### Definition 5

(Strict Step – Time Condition Shortest Path (SS–TCSP)) *This variant is the same as SSR – TCSW in that every edge used must move up in time. However, the difference in this variant is that the solution must be a path, not a walk (i.e. cannot traverse the same vertex twice, which implies no edge can be reused)*.

#### Definition 6

(Repay – Time Condition Shortest Walk (R–TCSW)) *Given the same inputs as TCSW, the problem becomes to find an a – b walk W from G*_1_ *to G_T_ where edges are paid for per use (rather than just once)*.

#### Definition 7

(Monotonic–Time Condition Shortest Walk (Mon–TCSW)) *This variant is similar to vanilla TCSW, except now* (*v_i_*, *w_j_*) *is a valid edge if v* ∈ *V_i_, w* ∈ *V_j_ and i* ≤ *j. We note in k — TCSW, we allow this jump to be up to step size k, but in this variant jumps can be arbitrarily large as long as they are monotonic*.

#### Definition 8

(Multi–k Time Condition Shortest Walk (Multi–k–TCSW)) *In this variant, we allow for a set of n demands 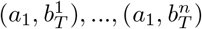. Our goal is to find a set of n walks starting from a singular source *a*_1_, and ending at 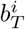. The cost paid is the sum of the weight of all edges used in one or more walks (each edge is only paid for at most once). This variant is most suitable for protein signalling cascades, whereby often times a signal begins at a singular receptor, and through a series of interactions, many downstream proteins are affected. Naturally k – TCSW is special case with only one demand*.

## Our Results

In this work, we build off the results of Wu et al. In particular we aim to show that similar to the Condition Shortest Path problems, extending the Condition setting to TCSW, a singular global shortest path problem with the *temporal walk constraint*, becomes hard to solve and hard to approximate to any non-trivial factor. We then relax the *temporal walk constraint*, and show a neat tradeoff between the temporal walk constraint and the hardness of approximation. Lastly, we present an ILP formulation for these problems, and demonstrate the ability to find optimal solutions to the generalized k-TCSW problem in feasible time on real world applications over the human Protein Protein Interaction Network (PPI).

We begin by proposing a rather simple algorithm for solving the *TCSW* problem for *T* ≤ 2.

### Theorem 1

*TCSW can be solved in polynomial for T* ≤ 2.

While this result is promising, the regime of polynomial time algorithms ends here.

### Theorem 2

*TCSW is hard to approximate to a factor of* 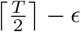 *for every fixed T* ≥ 3 *and ε* > 0.

Thus the best approximation ratio one can hope for is 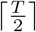. Such ratio can easily be achieved by considering the global network 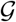 defined amongst all *G_i_* as follows: Create a network 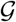 with node 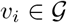 if *v_i_* ∈ *V_i_*.

An edge *e* = (*v_i_*, *w_j_*) exists in 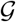 if *v_i_* ∈ *V_i_*, *w_j_* ∈ *V_j_*, *i* = *j* or *i* = *j*+1, and (*v_i_*, *w_j_*) ∈ *E*. In addition, we add zero weight edges for all nodes *v_i_*, *v*_*i*+1_ if *v_i_* ∈ *V_i_* and *v*_*i*+1_ ∈*V*_*i*+1_. Find the shortest path in this network from *a*_1_ to *b_T_* (thus ignoring the benefits of re-using an edge). An edge may be re-used once per alternate level, thus giving us a solution as bad as 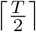 away from OPT.

By relaxing the *temporal walk constraint,* we are left with a more general result

### Theorem 3

*The general problem of k-TCSW is hard to approximate to a factor of* 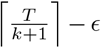 *for every fixed k* ≥ 1, *T* ≥ *k* + 2, and *ε* > 0.

Similar to vanilla TCSW, we can generate a global network 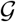 with node 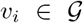 if *v_i_* ∈ *V_i_*. An edge *e* = (*v_i_*, *w_j_*) exists in 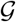 if *v_i_* ∈ *V_i_*, *w_j_* ∈ *V_j_*, *i* ≤ *j* ≤ *j*+*k*, and (*v_i_*, *w_j_*) ∈ *E*. In addition, we add zero weight edges for all self nodes up to *k* timepoints away (rather than 1). Similarly find the shortest path in this network from *a*_1_ to *b*_T_. In this instance, an edge may be reused once per *k* + 1 level, thus giving us a solution as bad as 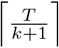 away from OPT.

Although these results are rather pessimistic in the theoretical abstract we provide a more consoling view by formulating an integer linear program for the general TCSW problem. We then show in experiments on real-world inputs derived from the human PPI network, the ILPs are capable of finding optimal solutions in a reasonable amount of time.

## Hardness of Time Condition Shortest Walk

### Theorem 1

*TCSW can be solved in polynomial for T* ≤ 2.

*Proof* Note that when *T* = 1 we’re simply left with the vanilla shortest path problem, which can easily be solved by algorithms such as Djikstra’s or Thorup’s Algorithm.

Now let us consider the case for *T* = 2. We argue that the optimal walk is in fact a path, and thus applying shortest path on the global network 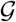 will provide the optimal solution.

Consider the global network 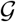 as defined earlier in Definition 1. Assume for the sake of contradiction that there exists a walk *W* that has a lower cost than the shortest path *P* from *a*_1_ to *b*_2_ in 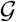. Due to our assumption and by definition of 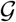, this walk cannot be a simple path. Therefore that implies there exists at least one vertex *v* ∈ *V* that was traversed twice in this walk, once from (*v_i_*, *w_j_*) and once from (*v_i′_*, *x_j′_*). Now consider the walk *W*′ formed by concatenating this portion of the walk and simply taking the edge (*v_i_*, *x_j′_*). As we only have two time points, this is a still a valid traversal and satisfies the *temporal walk constraint*. We can therefore apply this concatenation process until every vertex is traversed only once, thus forming a simple path *P*′, which has less than or equal cost to *W*. Therefore we arrive at a contradiction, as the optimal walk from *a* to b is in fact a path.

### Theorem 2

*TCSW is hard to approximate to a factor of* 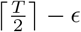 *for every fixed T* ≥ 3 *and ε* > 0.

We approach this proof via approximation preserving reduction from Condition Shortest Path to TCSW. Recall that in the Condition Shortest Path problem, we are given networks *G*_1_,…,*G_C_* and a source target pair (*a, b*). The goal is to find a subnetwork *H* ⊆ *G* = (∪_*c*_G_*c*_ of minimal cost such that there exists a path from *a* to *b* in H amongst vertices that are active in *G_c_*, for all *c*.

Therefore, given an instance of CSP (*G*_1_,…, *G_C_*, (*a, b*)), with underlying network *G* = (*V, E*) the reduction works as follows:

- Construct an instance of TCSW with 2C+1 networks 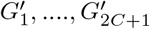. Let the odd networks in our constructed instance of TCSW 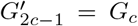 for *c* ∈ [*C*].
- For even instances *G*_2c_ for *c* ∈ [*C*], let *G*_2c_ be a network with exactly one node *t*\*.
- In addition to the edges *E* from our CSP instance, add two directed edges to our global edge set: (*b*, *t*\*) and (*t*\*, *a*), which we call our transition edges.
- Our singular demand is to find a walk from *a*_1_ to *b*_2*C*+1_
- The network *H* ⊆ *G* that satisfies CSP is formed by taking the union of edges traversed in the TCSW instance solution *W* minus the transition edges (*b*, *t*\*) and (*t*\*, *a*)

We first note that by construction, the only way to go from *a*_1_ to *b*_2*C*+1_ is to first go from *a*_1_ to *b*_1_, then take the transition edges from *b*_1_ to 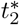 and 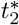 to *a*_3_, and then repeat. Therefore *W* contains a path from *a_c_* to *b_c_* that only goes through edges in *G_c_* for all *c* ∈ [*C*], and therefore the union of edges *e* ∈ *W* = *H* forms a valid solution for our CSP instance.

Now assume for the sake of contradiction that there exists a solution 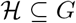 of lower overall cost than the network *H* returned from our TCSW reduction. For each condition *c* ∈ [*C*], consider an *a* – *b* path going through *H* using only edges in *G_c_*. Let the set of edges used in such path be called *P_c_*. We construct a walk 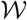 as follows: 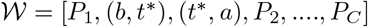. This forms a valid walk in our TCSW instance, and has lower cost than *W*, as edge weights are only paid for once in both the CSP instance and the TCSW instance. This leads to a contradiction that *W* was our optimal walk.

Therefore, all CSP instances can be solved via a reduction to TCSW. Naturally, this leads us to the inapproximability of TCSW. Namely, CSP cannot be approximated to a factor of *C* – *ε* for all fixed *C* ≥ 2 and *≥* > 0. Therefore, by extension TCSW cannot be approximated to a factor 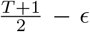 for all fixed odd *T* ≥ 3 and *ε* > 0, as we have 2*C* + 1 temporal conditions in our TCSW instance for a CSP instance of C conditions.

We note one caveat, which is that we’re always reducing to an odd number of time conditions. It is straight forward to see that we could have equivalently reduced to an even number of time conditions by adding an extra network *G*_2*C*+2_ with just the node *b*, and instead solving for the walk from *a*_1_ to *b*_2*C*+2_, thus proving that even instances are inapproximable to a factor of 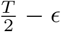 for all fixed even *T* ≥ 4 and *ε* > 0. As a result, we can make a more general statement from these two that TCSW is inapproximable to a factor of 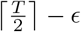

### Theorem 3

*The general problem of k-TCSW is hard to approximate to a factor of* 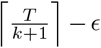 *for every fixed k* ≥ 1, *T* ≥ *k* + 2, *and ε* > 0.

This proof follows directly from Theorem 2. The reduction from CSP to k-TCSW follows the same flavor as the one above with the following key difference:

- Rather than constructing 2*C* + 1 networks, we construct (*k* + 1)*C* + 1 networks. We let every (*c* – 1)(*k* + 1) network 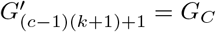.
- In addition, we let 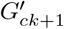 be a network with exactly one node t*
- All other networks 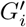 are empty
- We maintain the edge set formed previously, with the same two transition edges (*b*,*t*\*) and (*t*\*, *a*).
- Our singular demand is to find a walk from *a*_1_ to *b*_*C*(*k*+1)+1_
- The network *H* ⊆ *G* that satisfies CSP is formed by taking the union of edges traversed in the TCSW instance solution *W* minus the transition edges (*b*, *t*\*) and (*t*\*, *a*)

We note that the exact same arguments work for why *H* is a valid and optimal solution for the CSP instance. One caveat that the reader may notice, similar to the odd even bifurcation in the prior section is that we always reduce CSP to a k-TCSW instance in the form (*k* + 1)(*C* – 1) + 1. However, with the same idea of adding dummy networks at the end of our k-TCSW, we encompass all other instances of k-TCSW. As CSP is hard with *C* = 2, TCSW is NP-hard for all instances (*k*+1)(2–1)+1 = *k* + 2. In addition, k-TCSW is hard to approximate to factor 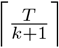 by similar argument.

### Building the protein-protein interaction network

In order to construct a human protein protein interaction (PPI) network network for our simulations, we collected data to construct a weighted directed network from four sources. Our largest dataset came from InWeb [24], where protein-protein interactions were treated as bidirectional edges from the proteins used. Edge weights were set as the negative log confidence score collected. Similarly, PPIs from the Human Protein Reference Data (HPRD)[25] were treated as bidirectional, but assigned the minimum nonzero confidence values by default. For directed edges, collected from highly curated data-sets, we used Phosphosite[26] and NetPath[27], and assigned edges sourced from both our maximal confidence value

### Solving k-TCSW to optimality

We can derive a natural linear program for k-TCSW in terms of network flows, by demanding a unit of flow from timepoint 1 for source *s* to arrive at timepoint *T* for target *t,* while maintaining the temporal walk constraint. We present the specific formulation in Figure 3.

**Figure 1.**
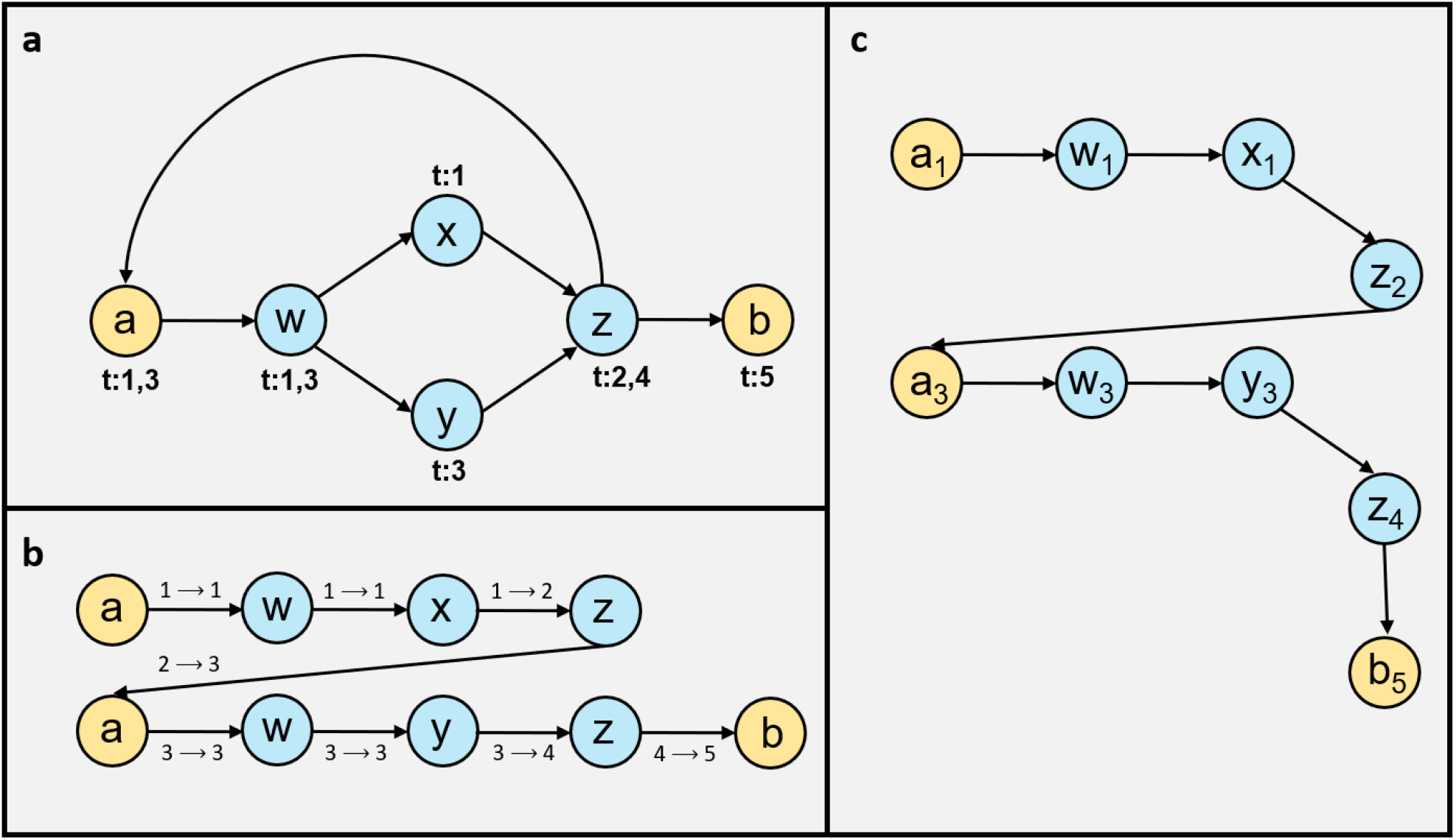
Example TCSW instance, with active time points per node labeled and source *a* and target *b* highlighted in yellow. In particular, we note that z is inactive at timepoint 3, and therefore the only possible solution is to transition from z in timepoint 2 to a in timepoint 3, ultimately going through w y and z again to get to b at timepoint 5. **(b)** Solution to the example instance. **(c)** Solution to the example instance based on the alternate formulation of 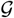 presented in Definition 1

**Figure 2.**
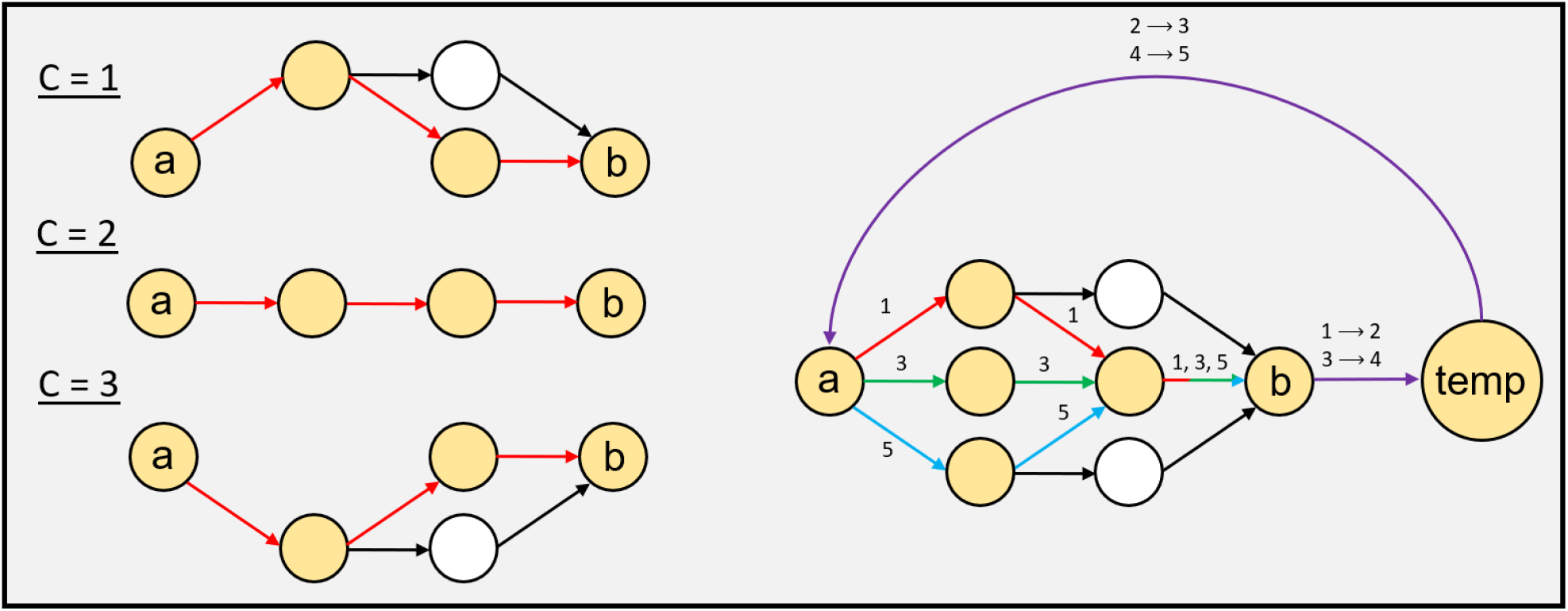
Example reduction from a Condition Shortest Path instance (left) with 3 conditions to an instance of TCSW with 5 time points (right). In the TCSW instance, red edges are traversed in timepoint 1, green edges in timepoints 3, blue edges in timepoint 5, and purple edges are traversed from timepoint *i* to timepoint *i* + 1.

**Figure 3.**
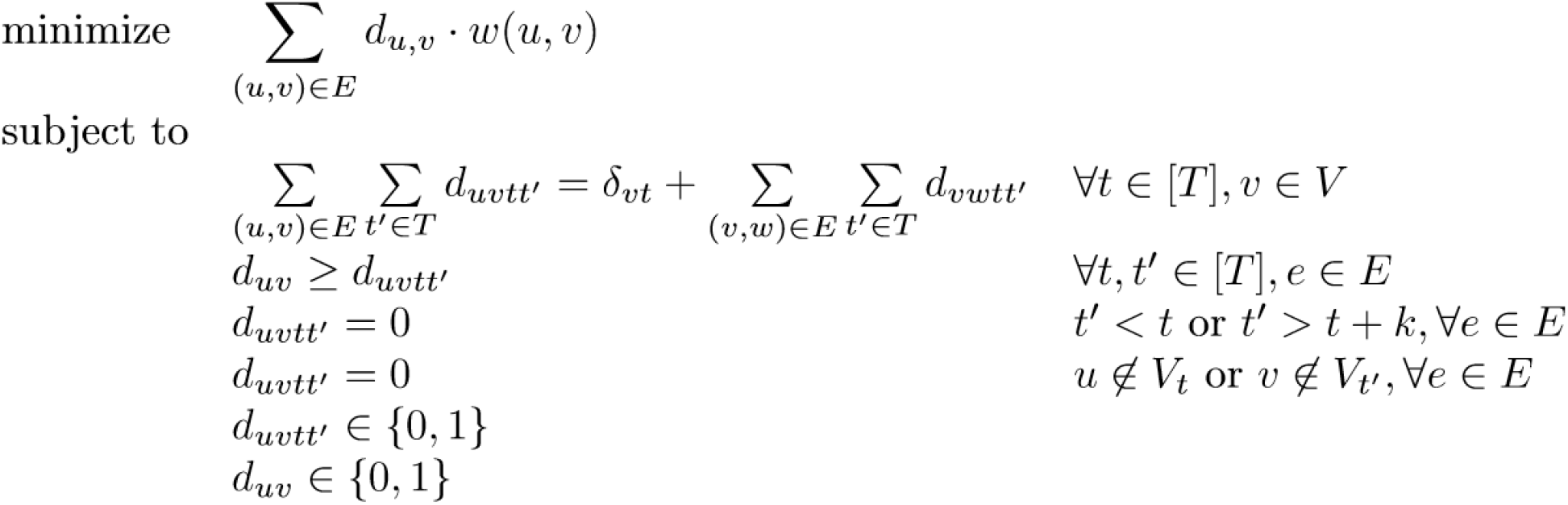
Integer linear program for k-Time Conditioned Temporal Walk. *δ_vt_* = 1 for *v* at time 1 if *v* is the source *s*, –1 if *v* is target *t* at time *T*, 0 otherwise. Each variable *d_uvtt′_* denotes the flow through edge (*u, v*) from time *t* to time *t′*; each variable *d_uv_* denotes whether (*u, v*) is ultimately in the chosen walk solution;. The first constraint enforces flow conservation by demanding 0 flow through all nodes except the source *s* and target *t*. The second constraint ensures that if an edge is used at any condition, it is chosen as part of the solution. The third constraint ensures that a jump of no larger than *k* is taken by forcing 0 flow through edges of greater time length. The fourth constraint ensures that both ends *u* and *v* exist in *V_t_* and *V_t′_* respectively.

### Performance analysis of integer linear programming

Given the protein-protein interaction network *G*, we sample an instance of *k*-TCSW by sampling a source node *a* ∈ *V*_1_ and target node *b* ∈ *V_T_* such that there exists a walk from *a* to *b* which satisfies the *k* temporal walk constraint. All other nodes exist in *G_t_* with probability *p*.

Using a workstation running an Intel Xeon E5-2690 processor and 250GB of RAM, **optimal solutions** to instances of modest size (generated using the procedure just described) were within reach:

**Table 1.** ILP solve times for some random instances generated by our random model using the Gurobi Python Solver package[28].

| $k$ | $T$ | $p$ | Time to solve |
| --- | --- | --- | --- |
| 1 | 1 | 0.25 | 40s ± 5s |
| 1 | 1 | 0.75 | 40s ± 5s |
| 1 | 3 | 0.25 | 4m 30s ± 1m |
| 1 | 3 | 0.75 | 20m ± 4m |
| 1 | 5 | 0.25 | 14m ± 6m |
| 1 | 5 | 0.75 | 24m ± 30s |
| 2 | 5 | 0.25 | 20m ± 6m 30s |
| 2 | 5 | 0.75 | 25m ± 30s |
| 1 | 10 | 0.50 | 31m ± 1m 30s |

We note that runtime seems to largely depend on the number of time conditions *T*, with some additional dependence on *p*, which loosely measures the size of our frames *G_t_*. Through these simulations we present a model which is applicable to real world biological instances, and can solve for optimal solutions in a feasible amount of time.

## Conclusion and discussion

In this manuscript we introduced the Time Condition Shortest Walk (TCSW) problem, in which the goal was to find an *a* – *b* path beginning in an initial time conditioned frame *G*_1_, and ending in *G_T_*, with jumps of length at most one. Unlike the shortest path problem, which is tractable, we demonstrated via approximation preserving reduction to Condition Shortest Path (CSP) that TCSW is hard to approximate within a factor 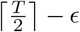 for *T* ≥ 3 and *ε* > 0. We then expanded this problem to a broader definition, k-TCSW, allowing for jumps of up to size *k* in this *a* – *b* path. We demonstrated a direct inverse relation between the jump size *k*, and best approximation ratio achievable, with *k–TCSW* being hard to approximate to 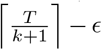 for *T* ≥ 3 and *ε* > 0 *k* ≥ 1. Lastly, we developed an integer linear program modeled on network flows, and applied it to solve for exact solutions over simulated instances on the human protein-protein interaction network, demonstrating feasibility runtimes for real-world instances.

In this work we also briefly explored a variety of alternative formulations for this problems, some being tractable, and others intractable. We believe a natural extension of this work would be to define the time conditioned frames over a series of variable edges, or both variable edges and vertices. Some modifications would have to be made to account for the ability to traverse self edges. Lastly, we believe given the feasibility demonstrated by our simulations, the next step would be to apply this method to cell signalling data spanning multiple time points.

## Analysis of problem variants

### Proposition 1

*Strict Step Repay - Time Condition Shortest Walk (SSR-TCSW) can be solved in polynomial time*

*Proof* We provide a simple algorithm to solve the problem in *O*(|*T*||*E*| + |*T*||*V*|*log*(|*T*||*V*|))

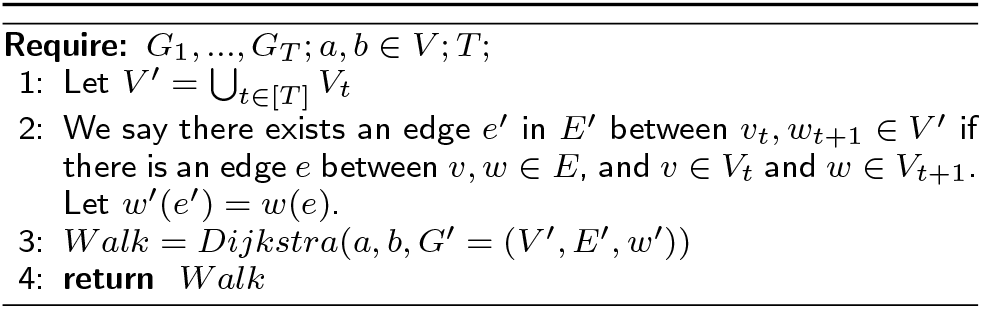

Note that this is just a simple modification of the shortest walk problem where we must use exactly *T* – 1 edges. By construction each edge moves us up exactly one time point. The only difference is that we cannot pass through certain vertices at certain time points, which we account for by not including the corresponding vertices in our modified network *G*′.

### Proposition 2

*Strict Step - Time Condition Shortest Path (SS-TCSP) is NP-Hard*

*Proof* We will show a simple reduction from Hamiltonian Path to SS-TCSP. Consider an instance of Hamiltonian Path, where given a graph *G* = (*V*, *E*), we are asked to find a simple path *P* s.t. *P* visits all vertices of G.

Let |*V*| = *n*. We now generate a new instance of SS-TCSP. First initialize *G*_2_,…,*G*_*n*+1_ = *G*. Initialize *G*_1_ with a singular source node *s* and *G*_*n*+2_ with a target node *t*. Define *T* = *n* + 2. Initialize *E*′ = *E*, and add to *E*′ edges (*s, v*) and (*v, t*) for *v* ∈ *V*

Now consider the instance *I* = {*G*_1_,…*G*_*n*+2_, (*s, t*), *T*}. We note that during time 2,..,—V—+1 the path must stay in the original *G*, and based on the definition of Simple Path, we are not allowed to visit a node more than once. Therefore, if a solution exists to *I*, it must go through each node in *G* exactly once, which can only be the case if and only if there is a Hamiltonian Path in *G*. Therefore, any instance of Hamiltonian Path can be reduced to an instance of TCP, which implies TCP is NP-Hard.

### Proposition 3

*Repay-Time Condition Shortest Walk (Mon-TCSW) can be solved in polynomial time*

*Proof* We note that the solution to this problem is rather trivial. Recall the network 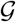 from Definition 1. Run Djikstra’s starting from *a*_1_ to find the shortest path to *b_T_*. By construction of 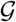 the walk satisfies the temporal walk constraint. In addition, as edges are paid for per use this will find the optimal walk from *a* to *b* spanning *G*_1_,…,*G_T_*.

### Proposition 4

*Monotonic-Time Condition Shortest Walk (Mon-TCSW) can be solved in polynomial time*

### Lemma 1

*An edge e* ∈ *E is traversed more than once in W only if there is a cycle in W*

*Proof* Let *e* ={*v, w*}. If e appears in *W* more than once, this implies that *v* was visited more than once, which implies there is a cycle in *W*.

### Lemma 2

*There exists an optimal walk W for Mon-TCSW that is a **Simple Path***.

*Proof* Assume for the sake of contradiction that there exists an walk *W′* that contains cycles that is better than *W*, our best Simple Path.

Let *v* be a node visited twice. This implies that *W*′ = {…, {*v*_1_, *w*},…{*v*2, *y*},…}, *w, y* ∈ *V*. However as *y* appears temporally after *v*_2_ in the path, it appears temporally after *v*_1_, which implies we can concatenate the cycle and get *W*\* ={…, {*v*_1_, *y*},…} which has equal or lower cost than *W*′ as all edge weights are positive. By applying this argument inductively, we arrive at a Simple Path *P* with no cycles that has lower cost than *W*′. Contradiction.

This implies the optimal walk *W* is a **Simple Path**.

### Lemma 3

*The optimal path does not visit each edge more than once once. Alternatively, it is enough to consider* min Σ_*e*∈*w*_^*w*(*e*)^

*Proof* By Lemma 2, the optimal walk W is a simple path. By Lemma 1, no edge is visited more than once.

Therefore, we simply suggest a modified version of the algorithm in for SSR-TCSW. Instead of simply introducing edges in SSR-TCSW from *v_t_* to *w*_*t*+1_, we introduce edges from *v_t_* to w_t′_ ∀*t*′ ≥ *t* as long as (*v, w*) ∈ *E*. Therefore, this gives us a simple algorithm to solve the problem in *O*(|*T*|^2^|*E*|+|*T*||*V*|*log*(|*T*||*V*|)).

## Formal Definition of TCSW and its variants

### Definition 9

(Time Condition Shortest Walk (TCSW)) *Consider the following inputs*:

1. *A global weighted network* 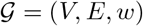
2. *A temporal activity function ρ*: (*V*, {1 .. *T*}) → {0, 1}, *indicating whether a node v* ∈ *V was active during time point* t ∈ {1… *T*}
3. *A pair of nodes* (*a, b*) ∈ *V*

We define 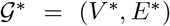 as the *ρ-unwinding* of 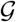. In this graph, the set of nodes *V*\* includes multiple copies of every node in *V*, with one copy for each time point in which it was active. Formally, denote the copy of *u* ∈ *V* at time t as *u_t_* and let *V*_t_ ={*u_t_* s.t. *u* ∈ *V* ∧ *ρ^t^*(*u*) = 1}. We then define *V*\* = (∪_*t*_*V_t_*. Similarly, we define *E*\* as the set of edges that connect node instances from the same time points or from adjacent time points. Formally, let *E_t_* ={< *u_t_*, *v*_*t*+*i*_ > *s.t*. (*u* = *v* ∨ < *u, v* > ∈ *E*) ∧ (*ℓ* ≤ *i* ≤ *k*)}

Notably, any path *P* ∈ *E*\* would be time consistent - namely it will only include transitions between contemporary nodes or between nodes at adjacent time points, moving up in time. For a path *P* ∈ *E*\* we denote by *c*(*P*) the cost with edge repayment per use and *c*\*(*P*) the cost without repayment per use (edges traversed at two different time points are considered the same edge). In the TCSW problem, we have a **ρ*-unwinding* of 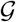 with *ℓ* = 0, *k* =1 and optimize for *c*\*(*P*). In the SSR-TCSW problem, we use *ℓ* = 1, *k* = 1 and optimize for *c*\*(*P*). In the Mon-TCSW we use *ℓ* = 0, *k* = *T* — 1 and optimize for *c*(*P*).

## List of Abbreviations

TCSW: Time Condition Shortest Walk
k-TCSW: k–Time Condition Shortest Walk
SSR-TCSW: Strict Step Repay–Time Condition Shortest Walk
SS-TCSP: Strict Step – Time Condition Shortest Path
R–TCSW: Repay–Time Condition Shortest Walk
Mon–TCSW: Monotonic–Time Condition Shortest Walk
Multi–k–TCSW: Multi–k Time Condition Shortest Walk
PPI: protein-protein interaction

## Declarations

### Ethics approval and consent to participate

Not applicable

### Consent for publication

Not applicable

### Availability of data and material

Our simulation code can be found at the following URL: https://github.com/YosefLab/temporal_condition_shortest_walk

### Author’s contributions

All authors conceived and designed the study. AK derived the main hardness results, the ILP formulation, and the benchmarking. NY was the PI and oversaw the project.

### Competing interests

The authors declare that they have no competing interests.

### Funding

This work was partially supported by the National Science Foundation Graduate Research Fellowship Program award DGE 1106400, NIH grants U01HG007910 and U01MH105979, and the U.S.-Israel Binational Science Foundation.

## Notes

### Competing Interest Statement

The authors have declared no competing interest.

## References

1. Wu, C., Yosef, N., Thalhamer, T., Zhu, C., Xiao, S., Kishi, Y., Regev, A., Kuchroo, V.K.: Induction of pathogenic th17 cells by inducible salt-sensing kinase sgk1. Nature 496(7446), 513–517 (2013)

2. Huang, S.S., Fraenkel, E.: Integrating proteomic, transcriptional, and interactome data reveals hidden components of signaling and regulatory networks. Sci Signal 2(81), 40 (2009)

3. Signorini, L.F., Almozlino, T., Sharan, R.: Anat 3.0: a framework for elucidating functional protein subnetworks using graph-theoretic and machine learning approaches. BMC Bioinformatics 22(1), 526 (2021). doi:10.1186/s12859-021-04449-1

4. Rauti, R., Shahoha, M., Leichtmann-Bardoogo, Y., Nasser, R., Paz, E., Tamir, R., Miller, V., Babich, T., Shaked, K., Ehrlich, A., Ioannidis, K., Nahmias, Y., Sharan, R., Ashery, U., Maoz, B.M.: Effect of sars-cov-2 proteins on vascular permeability. eLife 10, 69314 (2021). doi:10.7554/eLife.69314

5. Garmaroudi, F.S., Marchant, D., Si, X., Khalili, A., Bashashati, A., Wong, B.W., Tabet, A., Ng, R.T., Murphy, K., Luo, H., Janes, K.A., McManus, B.M.: Pairwise network mechanisms in the host signaling response to coxsackievirus B3 infection. Proceedings of the National Academy of Sciences 107(39), 17053–17058 (2010). doi:10.1073/pnas.1006478107. https://www.pnas.org/content/107/39/17053.full.pdf

6. Scott, M.S., Perkins, T., Bunnell, S., Pepin, F., Thomas, D.Y., Hallett, M.: Identifying Regulatory Subnetworks for a Set of Genes. Molecular & Cellular Proteomics 4(5), 683–692 (2005)

7. Huang, S.-s.C., Fraenkel, E.: Integration of Proteomic, Transcriptional, and Interactome Data Reveals Hidden Signaling Components. Science signaling 2(81), 40 (2009)

8. Przytycka, T.M., Singh, M., Slonim, D.K.: Toward the dynamic interactome: it’s about time. Briefings in Bioinformatics 11(1), 15–29 (2010). doi:10.1093/bib/bbp057

9. Mertins, P., Przybylski, D., Yosef, N., Qiao, J., Clauser, K., Raychowdhury, R., Eisenhaure, T.M., Maritzen, T., Haucke, V., Satoh, T., Akira, S., Carr, S.A., Regev, A., Hacohen, N., Chevrier, N.: An Integrative Framework Reveals Signaling-to-Transcription Events in Toll-like Receptor Signaling. Cell Reports 19(13), 2853–2866 (XXXX). doi:10.1016/j.celrep.2017.06.016

10. Schiebinger, G., Shu, J., Tabaka, M., Cleary, B., Subramanian, V., Solomon, A., Gould, J., Liu, S., Lin, S., Berube, P., Lee, L., Chen, J., Brumbaugh, J., Rigollet, P., Hochedlinger, K., Jaenisch, R., Regev, A., Lander, E.S.: Optimal-transport analysis of single-cell gene expression identifies developmental trajectories in reprogramming. Cell 176(4), 928–94322 (2019). doi:10.1016/j.cell.2019.01.006

11. Holme, P.: Modern temporal network theory: A colloquium. The European Physical Journal B 88 (2015). doi:10.1140/epjb/e2015-60657-4

12. Holme, P., Saramäki, J.: Temporal networks. Physics Reports 519(3), 97–125 (2012). doi:10.1016/j.physrep.2012.03.001. Temporal Networks

13. Masuda, N., Lambiotte, R.: A Guide to Temporal Networks, (2016). doi:10.1142/q0268

14. Wu, H., Cheng, J., Huang, S., Ke, Y., Lu, Y., Xu, Y.: Path problems in temporal graphs. Proc. VLDB Endow. 7(9), 721–732 (2014). doi:10.14778/2732939.2732945

15. Wang, S., Lin, W., Yang, Y., Xiao, X., Zhou, S.: Efficient route planning on public transportation networks: A labelling approach. In: Proceedings of the 2015 ACM SIGMOD International Conference on Management of Data. SIGMOD ‘15, pp. 967–982. Association for Computing Machinery, New York, NY, USA (2015). doi:10.1145/2723372.2749456. https://doi.org/10.1145/2723372.2749456

16. Wu, H., Huang, Y., Cheng, J., Li, J., Ke, Y.: Reachability and time-based path queries in temporal graphs. In: 2016 IEEE 32nd International Conference on Data Engineering (ICDE), pp. 145–156 (2016). doi:10.1109/ICDE.2016.7498236

17. Nicosia, V., Tang, J., Musolesi, M., Russo, G., Mascolo, C., Latora, V.: Components in time-varying graphs. Chaos: An Interdisciplinary Journal of Nonlinear Science 22(2), 023101 (2012). doi:10.1063/1.3697996. https://doi.org/10.1063/1.3697996

18. Huang, S., Fu, A.W.-C., Liu, R.: Minimum spanning trees in temporal graphs. In: Proceedings of the 2015 ACM SIGMOD International Conference on Management of Data. SIGMOD ‘15, pp. 419–430. Association for Computing Machinery, New York, NY, USA (2015). doi:10.1145/2723372.2723717. https://doi.org/10.1145/2723372.2723717

19. Ikuta, T., Akiba, T.: Integer programming approach for directed minimum spanning tree problem on temporal graphs. In: Proceedings of the 1st ACM SIGMOD Workshop on Network Data Analytics. NDA ‘16. Association for Computing Machinery, New York, NY, USA (2016). doi:10.1145/2980523.2980528. https://doi.org/10.1145/2980523.2980528

20. Rozenshtein, P., Gionis, A., Prakash, B.A., Vreeken, J.: Reconstructing an epidemic over time. In: Proceedings of the 22nd ACM SIGKDD International Conference on Knowledge Discovery and Data Mining. KDD ‘16, pp. 1835–1844. Association for Computing Machinery, New York, NY, USA (2016). doi:10.1145/2939672.2939865. https://doi.org/10.1145/2939672.2939865

21. Charikar, M., Chekuri, C., Cheung, T.-y., Dai, Z., Goel, A., Guha, S., Li, M.: Approximation algorithms for directed steiner problems. Journal of Algorithms 33(1), 73–91 (1999). doi:10.1006/jagm.1999.1042

22. Byrka, J., Grandoni, F., Rothvoß, T., Sanita, L.: An improved lp-based approximation for steiner tree. In: Proceedings of the Forty-second ACM Symposium on Theory of Computing, pp. 583–592 (2010). ACM

23. Wu, J., Khodaverdian, A., Weitz, B., Yosef, N.: Connectivity problems on heterogeneous graphs. Algorithms for Molecular Biology 14(1), 5 (2019). doi:10.1186/s13015-019-0141-z

24. Li, T., Wernersson, R., Hansen, R.B., Horn, H., Mercer, J., Slodkowicz, G., Workman, C.T., Rigina, O., Rapacki, K., Stærfeldt, H.H., Brunak, S., Jensen, T.S., Lage, K.: A scored human protein-protein interaction network to catalyze genomic interpretation. Nature Methods 14, 61 (2016)

25. Keshava Prasad, T.S., Goel, R., Kandasamy, K., Keerthikumar, S., Kumar, S., Mathivanan, S., Telikicherla, D., Raju, R., Shafreen, B., Venugopal, A., Balakrishnan, L., Marimuthu, A., Banerjee, S., Somanathan, D.S., Sebastian, A., Rani, S., Ray, S., Harrys Kishore, C.J., Kanth, S., Ahmed, M., Kashyap, M.K., Mohmood, R., Ramachandra, Y.L., Krishna, V., Rahiman, B.A., Mohan, S., Ranganathan, P., Ramabadran, S., Chaerkady, R., Pandey, A.: Human Protein Reference Database—2009 update. Nucleic Acids Research 37(suppl_1), 767–772 (2009). doi:10.1093/nar/gkn892

26. Hornbeck, P.V., Zhang, B., Murray, B., Kornhauser, J.M., Latham, V., Skrzypek, E.: Phosphositeplus, 2014: mutations, PTMs and recalibrations. Nucleic Acids Res 43(Database issue), 512–520 (2015). doi:10.1093/nar/gku1267

27. Kandasamy, K., Mohan, S.S., Raju, R., Keerthikumar, S., Kumar, G.S.S., Venugopal, A.K., Telikicherla, D., Navarro, J.D., Mathivanan, S., Pecquet, C., Gollapudi, S.K., Tattikota, S.G., Mohan, S., Padhukasahasram, H., Subbannayya, Y., Goel, R., Jacob, H.K., Zhong, J., Sekhar, R., Nanjappa, V., Balakrishnan, L., Subbaiah, R., Ramachandra, Y., Rahiman, B.A., Prasad, T.K., Lin, J.-X., Houtman, J.C., Desiderio, S., Renauld, J.-C., Constantinescu, S.N., Ohara, O., Hirano, T., Kubo, M., Singh, S., Khatri, P., Draghici, S., Bader, G.D., Sander, C., Leonard, W.J., Pandey, A.: Netpath: a public resource of curated signal transduction pathways. Genome Biology 11(1), 3 (2010). doi:10.1186/gb-2010-11-1-r3

28. Gurobi Optimization, L.: Gurobi Optimizer Reference Manual (2018). http://www.gurobi.com

